# Bodily ownership of an independent supernumerary limb: An exploratory study

**DOI:** 10.1101/2021.09.13.459945

**Authors:** Kohei Umezawa, Yuta Suzuki, Gowrishankar Ganesh, Yoichi Miyawaki

## Abstract

Can our brain perceive a sense of ownership towards an *independent* supernumerary limb; one that can be moved independently of any other limb and provides its own independent movement feedback? Following the rubber-hand illusion experiment, a plethora of studies has shown that the human representation of ‘self’ is very plastic. But previous studies have almost exclusively investigated ownership towards ‘substitute’ artificial limbs, which are controlled by the movements of a real limb and/or limbs from which non-visual sensory feedback is provided on an existing limb. Here, to investigate whether the human brain can own an independent artificial limb, we first developed a novel *independent* robotic ‘sixth finger.’ We allowed participants to train using the finger and examined whether it induced changes in the body representation using behavioral as well as cognitive measures. Our results suggest that unlike a substituted artificial limb (like in the rubber hand experiment), it is more difficult for humans to perceive a sense of ownership towards an independent limb. However, ownership does seem possible, as we observed clear tendencies of changes in the body representation that correlated with the cognitive reports of the sense of ownership. Our results provide the first evidence to show that an independent supernumerary limb can be embodied by the human brain.

## Introduction

The representation of our body in the brain is plastic. Even relatively short multi-sensory (usually visuo-haptic) stimulations can induce an illusion of ownership, and embodiment in general, towards artificial limbs and bodies (Aymerich-Franch and Ganesh, 2016) including human-looking limbs: rubber hands (Botvinick and Cohen, 1998; Longo et al., 2008), virtual avatars (Slater et al., 2009, 2010; Hagiwara et al., 2020) as well as robotic limbs and bodies (Nishio et al., 2012; Alimardani et al., 2013; Aymerich-Franch et al., 2015, 2017, 2016; Aymerich-Franch et al., 2016; Suzuki et al., 2015). However, most previous studies on ownership have been conducted in the context of ‘limb substitution,’ where the rubber, virtual or robot limb replaces an already-existing limb. Even studies that have explored the sense of ownership towards additional limbs such as extra hands (Ehrsson, 2009), tail (Steptoe et al., 2013), arm (Iwasaki and Iwata 2018) or finger (Llorens et al., 2012; Keiliba et al., 2021) for example, have utilized non-independent limbs because the movement of these additional limbs were controlled by other existing limbs. It remains unclear whether the human brain can own an independent additional limb that can be controlled independently of any other limbs and provide independent non-visual feedback and how it affects the body representations in the brain.

Ownership of a limb is believed to require two conditions to be satisfied. First, a congruent multi-sensory stimulation, a condition that was the key discovery of the rubber hand illusion by Botvinick and Cohen (Botvinick and Cohen, 1998). It and subsequent studies by other researchers have shown that the vision of an artificial limb (like the rubber hand) being touched and the haptic perception of the same touch from the real limb that is touched simultaneously are required for the induction of the illusion of ownership toward the artificial limb. However, an independent additional limb, by definition, does not have a replacement limb for providing this feedback. Furthermore, it is believed that the illusion of ownership occurs only with artificial limbs that conform to the so-called ‘body model,’ a representation of the body in our brain (Tsakiris and Haggard, 2005). What the body model constitutes of, however, remains unclear. Multiple studies on this issue (Haans et al., 2008; Guterstam et al., 2013; Pavani and Zampini, 2007; Pavani et al., 2000; Austen et al., 2004; Ehrsson et al., 2004; Costantini and Haggard, 2007; Lloyd, 2007 for example) have discovered diverse results that are probably best explained by the functional body model hypothesis (Aymerich-Franch and Ganesh, 2016) that is the entity that a brain can embody is determined by whether it is recognized to sufficiently afford actions that the brain has learnt to associate with the limb it substitutes. This would however suggest that the ownership of a new independent additional limb, which is not being associated to any actions, would therefore be difficult for humans. Overall, the previous studies therefore suggest that bodily ownership of an independent limb should be difficult to achieve.

To verify whether bodily ownership is possible for an independent limb, in this exploratory study we developed a robotic independent ‘sixth finger’ system. Our setup attaches the robotic finger to the participant’s hand and is operated by a ‘null-space’ electromyography (EMG) control strategy that enables individuals to operate the robotic finger and sense its movement independently of any other fingers or body parts. We then let participants experience using the finger through a cued finger pressing task in two conditions: one when they had control of the robotic finger (controllable condition) and the other when they could not control the robotic finger (the random condition). Following the habitation, we evaluated whether and how the use of fingers in the two conditions affected the participant’s perception of their innate hand by using cognitive measures collected with questionnaires (to measure changes in perceived levels of ownership and agency towards the robotic finger as well as body image of their hand) and behavioral measures collected in two test experiments that measured changes in their arm reaching to test for possible changes in their ‘body schema’ (Cardinalli et al., 2009; Maravita and Iriki, 2004) and a finger localization test in the visual space (to test possible changes in their ‘body image’, Sposito et al. 2012).

## Results

### The independent robotic sixth finger system

Our developed robotic sixth finger system consists of a three-phalanged, one degree of freedom, plastic finger (6 cm in length) and a tactile stimulator in the form of a sliding plastic pin. The finger was actuated by a servo motor (ASV-15-MG, Asakusa Gear Co., Ltd.) (Figure 1A) that enabled the robotic finger to flex like a real human finger. The robotic finger was connected to the stimulator by a crank-pin link such that the flexing of the finger made the pin slide. The entire arrangement was designed to be strapped onto the hand of participants, near their left little finger, such that the tactile stimulator stimulated the side of their palm, below the little finger, when the sixth finger flexed (see Figure 1A). The robot finger was operated using the electromyography (EMG) recorded from four wrist/finger muscles (flexor carpi radialis muscle, flexor carpi ulnaris muscle, extensors carpi radialis muscle, and extensor digitorum muscle) in their left forearm. We used the Delsys Trigno Wireless EMG System (sampling rate, 2000 Hz) for the EMG recording. The recorded EMG signals were read into the Arduino Mega 2560, utilizing the Simulink Support Package for Arduino Hardware.

**Figure 1:**
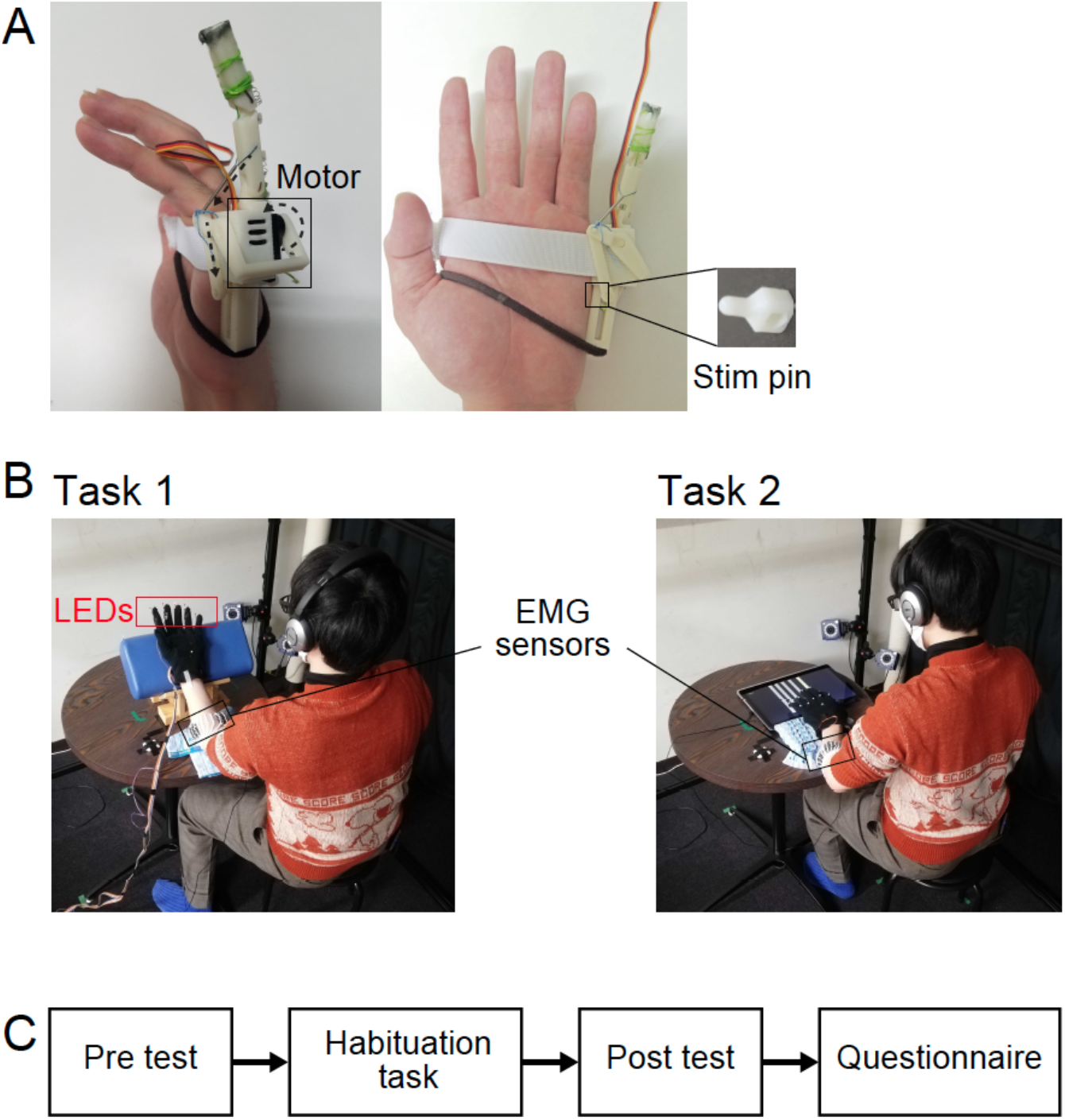
The sixth finger setup and experiment. (A) The 3D-printed robotic sixth finger actuated by a servo motor in the chassis. The entire arrangement is designed to be strapped onto the hand of participants, near their left little finger. (B) Schematic of the training task. In task 1, participants wore a glove with LEDs fixed on the tip of the glove fingers. They bent the finger corresponding to the led that lit up. In task 2, a tablet PC on the desk displayed a piano key under each of the five fingers (including the robot finger, and except the thumb). The participants pressed the key that lighted up. The task in both paradigms was aligned with the beats of a popular music that was played in the headphones of the participants. (C) The sequence of experiments.

Two salient features distinguish our sixth finger from most previous developments of supernumerary limbs-

1. Kinematically independent control: We developed an intuitive ‘null space’ control scheme for the robotic finger that enables the participant to move the robotic finger independently of, and simultaneously with if required, any other finger, and in fact any other limb on their body. Details of the control are provided in the methods section.
2. Independent haptic sensory feedback: The sliding pin stimulator incorporated with the finger provides haptic feedback of the movement of the robot finger on the side of the palm, and ensuring sensory independence to any other limbs of the body.

### Cognitive measures

To quantify the cognitive change in perception of the finger and hand following the training with the robotic finger, we asked participants to answer questionnaires about sense of agency (Q1, 3, 5), sense of ownership (Q4, 6), and body image (Q2, 7) by using seven-level Likert scale after habituation to the use of the robotic finger. Results showed significant differences in the questionnaire scores across the controllable and random conditions for the items related to the sense of agency (Wilcoxon sign-rank test, Q1: *T*(17) = 153, *p* < 0.0003, Q3: T(17) = 113.5, *p* < 0.0025, Q5: *T*(17)= 171, *p* < 0.0002), but not for the item related to sense of ownership (Q4: *T*(17) = 69, *p* > 0.05, Q6: *T*(17) = 62.5, *p* > 0.05) and to body image (Q2: *T*(17)= 93.5, *p* > 0.05, Q7: *T*(17) = 28, *p* > 0.05) (Figure 2).

**Figure 2:**
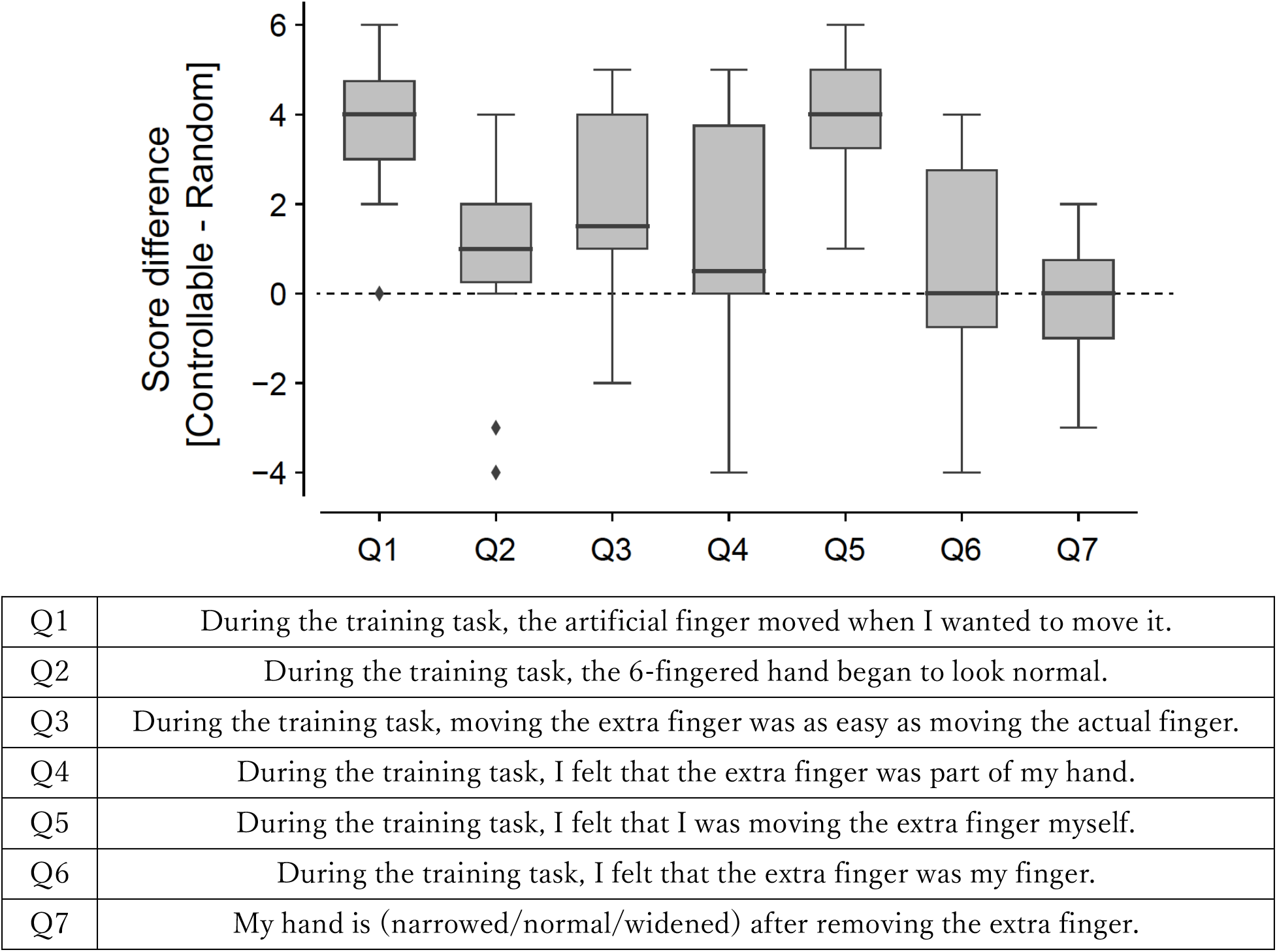
Results of subjective ratings of questionnaires. For each box plot, a thick line indicates the median, and the bottom and the top lines of the box show the first and the third quartiles, respectively. The whiskers extend to the minimum and maximum points of the data or to the points of 1.5 times of the interquartile range if the minimum and maximum points exceed them. The diamond markers showed the data point considered as outliers that exceed the whisker range.

### Behavioral tests

Together with the questionnaire, we also performed two behavioral tests to examine possible changes in body schema (reaching test) and body image (finger localization test) before and after the use of the robotic finger. In the reaching task, participants were instructed to reach with their index finger to a target line while avoiding an obstacle to impede the shortest path from the starting location of the index finger to the target location (see Figure 3A). Body schema is said to represent the internal representation of the body in the brain that is used for motor control (Cardinalli et al., 2009; Ganesh et al., 2014), and hence we expected an increased avoidance if there was a change in their body schema related to their hand, and the participants perceived their hand to be larger than before.

**Figure 3:**
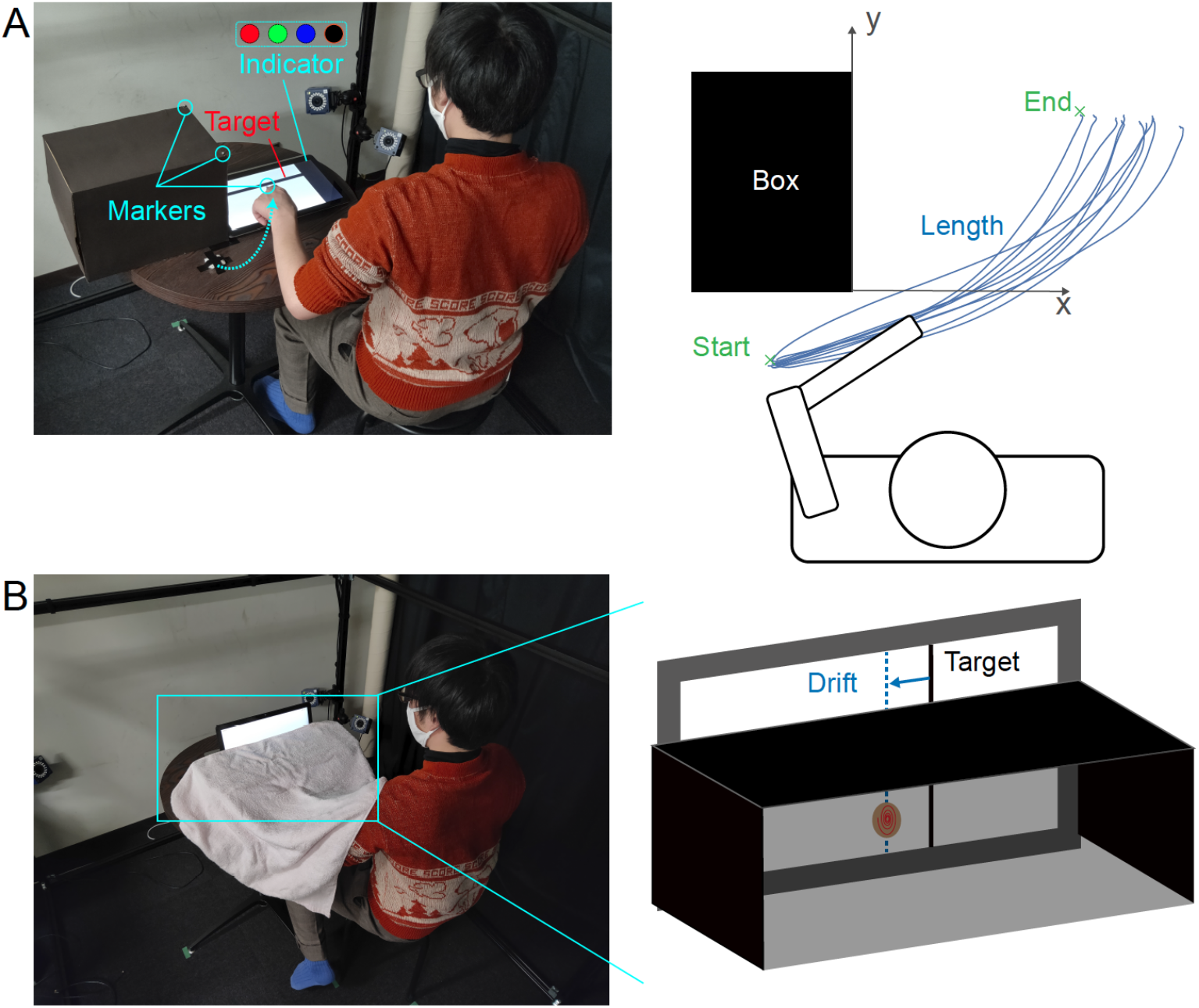
Measurement of body representation changes. (A) Schematic of the reaching test. A rectangular-shaped obstacle was placed in front of the participant’s left arm and a tablet PC was placed to the right of it. A target bar and an indicator were displayed on the screen. (B) Schematic of the finger localization test. An open box (without the front and back faces) and a PC standing behind it were placed in front of a participant. A target bar was displayed on the screen. The participants touched the target bar with either of the index or little finger.

In the finger localization test, participants were instructed to point to a line, visible above a box on the table, with their occluded index and little finger, below the box top (see Figure 3B). Body image is believed to be the representation of our body in the visual space, and we expected that any change in their body image related to their hand would lead to an error in the localization of the fingers by the participants, relative to the presented lines.

Results obtained in the behavioral test, however, didn’t show any significant change in any of these measures from pre-habituation to post-habituation, between the controllable and random conditions (Figure 4; Wilcoxon sign-rank test, 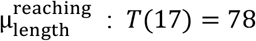, p = 0.766,, 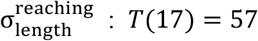, p = 0.229, 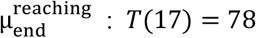, p = 0.766, 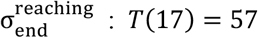, p = 0.229, 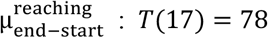, p = 0.766, 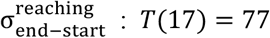, p = 0.734, 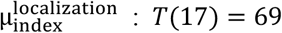, p = 0.495, 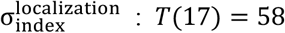, p = 0.246, 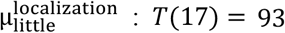, p = 0.766, 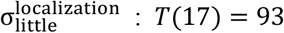, p = 0.766, 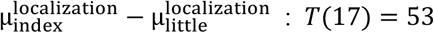, p = 0.167).

**Figure 4:**
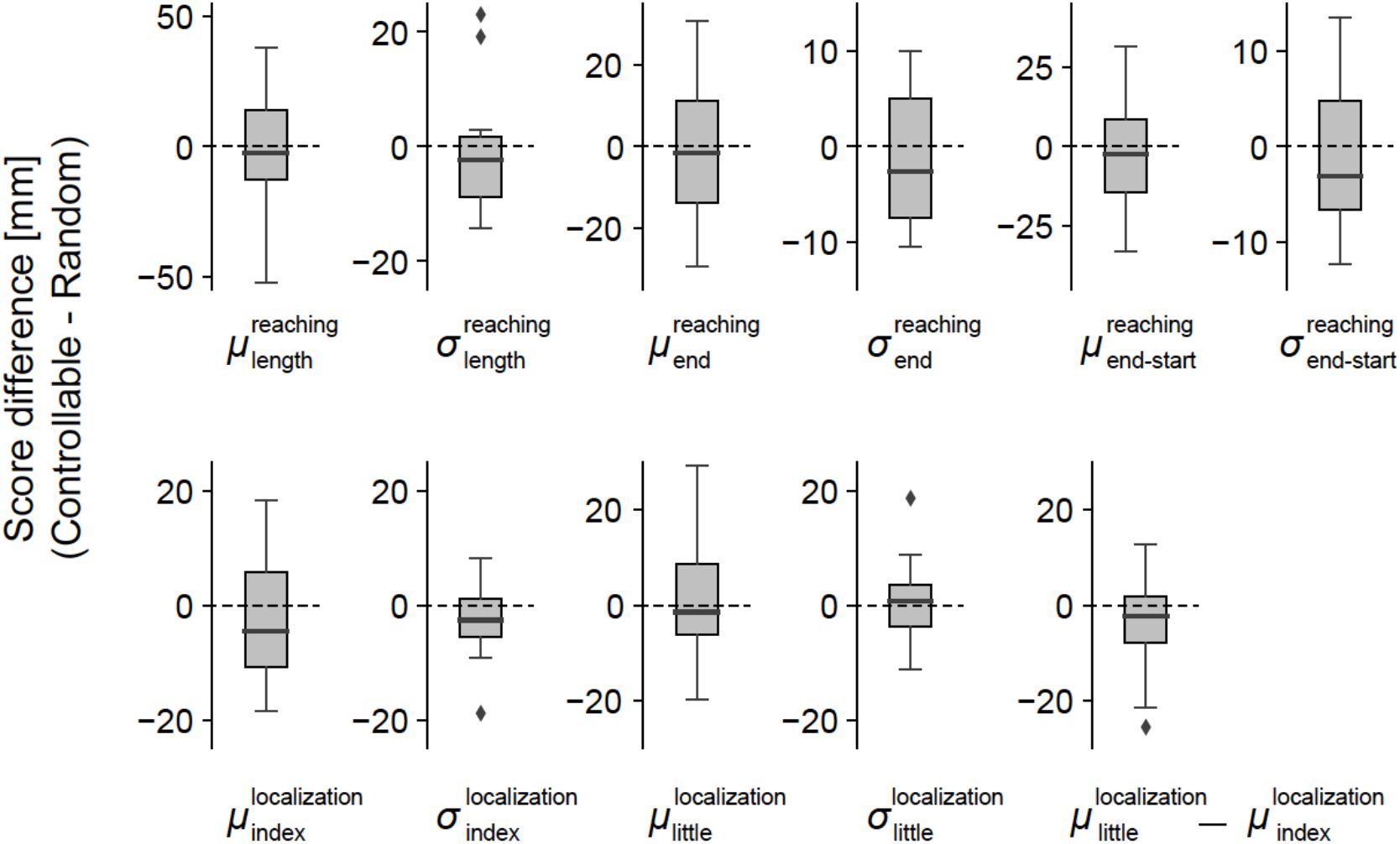
Results of behavioral measures (top, reaching test; bottom, localizaion test). The box plots showed the data as in the same style as Figure 2.

### Correlation between cognitive and behavioral measures

However, these insignificant differences in population levels may result from large variance in the behavioral measures and questionnaire scores across participants. Therefore, next we evaluated the correlation between the questionnaire score and behavioral measures across individuals. In a preliminary analysis, we observed a part of the data might serve as outliers (Smirnov-Grubbs test, α=0.01) that could distort estimates of the correlation between questionnaire score and behavioral measures if directly computed using all the data points. Therefore, instead of obtaining better estimates of correlation, we performed Bootstrap sampling to calculate the distribution of correlation values and determine the upper and lower limit within which 95% of the correlation values were distributed, which serve as a confidence interval of the correlation values.

Results showed significant correlations (as per the Bootstrap analysis) between the standard deviation in localization of the little finger 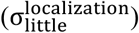 and the question related to whether the robotic finger was perceived as a part of their body (Q4, median R=0.568), as well as between 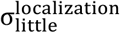 and the question of how much the robotic finger was perceived as their own finger (Q6, median R=0.571). These results suggest that the participant’s level of uncertainty in localizing little finger increased with the level of ownership they perceived cognitively toward the robotic finger.

## Discussion

In this exploratory study, we aimed to examine whether and to what extent humans can embody an ‘independent limb,’ a limb that humans can move independently of any other limbs in their body and from which they receive movement feedback independently of any other limbs. For this purpose, we developed a robotic finger setup that attaches to the hand of a human participant and can be moved by a muscle activity independently of that was used to move the other fingers of the hand, ensuring independence at the level of movement kinematics. Furthermore, we provided the participants with independent feedback corresponding to the finger movement via a tactile pin on the side of the palm.

In addition to a questionnaire to judge perceived changes in ownership, agency, and body image, we used two test experiments to evaluate possible changes in body representation after the use of the robotic finger. The first reaching test evaluated changes in the body schema, the representation of our body in internal coordinates that are used for motor control (Ganesh et al., 2014; Maravita and Iriki, 2004, Cardinali et al., 2009). Specifically, we evaluated changes in the representation of the width of the hand in the body schema. We expected the length of movement trajectory, and/or the distance between the start and end points of the reach to change in order to avoid the obstacle, in the case that the hand width is perceived to change. The second finger localization test was designed to analyze changes in the body image, the representation of our body in visual space (Iriki et al., 1996; Canzoneri et al., 2013). Here again, we expected the position of the index or little finger to change or become uncertain if the perceived visual hand width or perceived visual location of the little finger changed. We did not observe any change in the values of the questionnaire answers or the behavioral measures.

While the illusion of ownership towards a *replaced* rubber hand can be induced within a few minutes, our results suggest that the ownership of an *additional* limb is much tougher for the human brains. Although our EMG-finger interface is very intuitive and all our participants could operate the robotic finger almost immediately, participant’s reports of ownership did not differ between the controllable and random conditions even after 30 minutes of habituation. This difficulty in perceiving ownership agrees with the functional body model hypothesis (Aymerich-Franch and Ganesh, 2016). However, we observed clear tendencies of changes in body representation, shown as a clear correlation between the uncertainty in the position of the little finger and two questions that represent the ownership felt by the participants (Figure 5). These results give probably the first behavioral evidence to suggest that the human body can embody a truly independent supernumerary limb.

**Figure 5:**
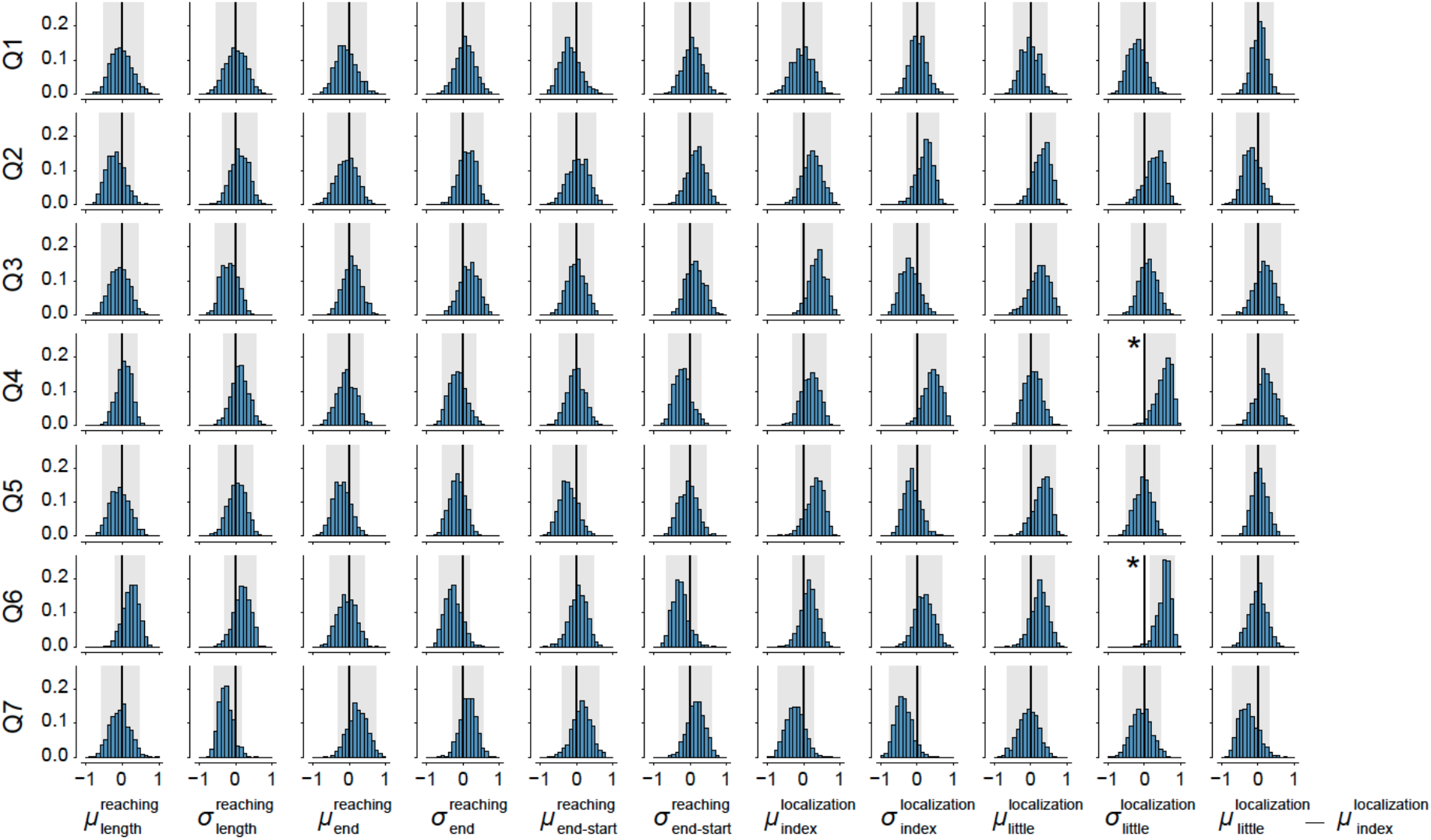
Bootstrap distributions of correlation values between differences across controllable and random conditions for questionnaire scores and post- and pre-habituation difference in behavioral measures. The black thick lines indicate zero correlation, and the gray shades indicate the range of correlation within which 95% of bootstrapped correlation values distributed for each combination of questions and behavior measures. The asterisks indicate correlation pairs whose 95% bootstrapped correlation distribution is not overlapped with zero correlation.

But is the robotic finger in fact treated as a tool rather than an additional limb? Previous studies have shown the human can embody tools into their body. Tools are additional to the human body by definition and are also known to change the body schema (Maravita and Iriki, 2004; Cardinali et al., 2009) and body image of the user (Sposito et al., 2012). However, one critical aspect that tool embodiment differs from limbs is in the lack of ownership, which is prominent only with limbs. And hence the correlated changes we observed between the behavior and ownership measure here suggest that our robotic finger tended to be considered as an additional limb by the participants.

In this study we could not clarify the role of haptic feedback in the embodiment. Our initial plan was to investigate also the embodiment differences between these groups. However, at least for the behavioral measures we considered in this study, we did not find a significant difference between the two groups. This may be due to two reasons. First, the fact that the feedback pin in the current design may not be conspicuous enough. Second, participants reported some haptic sensation from the robotic finger due to its movement dynamics, even in the absence. Further studies are required to clarify the effects of feedback on the embodiment of additional limbs.

## Methods

### Participants

Eighteen male participants (mean age 26.6 years, range 20 – 39 years, all right-handed) with normal or corrected to normal vision participated in this study. Each participant gave written informed consent before participating. The procedure was approved by the institutional review board of The University of Electro-Communications. Four participants who did not complete the experiments due to scheduling issues (one participant) or technical issues with the robot (three participants) are not included in the eighteen.

### Independent control of the robotic sixth finger

Human joints are actuated by a redundant set of muscles and antagonist muscles. The difference between the antagonist *flexor* and *extensor* muscles provides the torque to actuate a joint while the *co-activation*, or the common part of their activations, lets us increase the impedance of the joints irrespective of the applied torque. In this study, it was critical for us that the participants moved their six fingers (the five real and one robot finger) independently. We achieved this by isolating and utilizing the co-activation component of the muscle activations, which did not contribute to the motion of any of the real fingers, to actuate the robotic finger. The co-activation component in the muscle was isolated as follows.

Participants in our experiment started with a calibration session in which they were presented with a sinusoidal moving cursor and asked to repeatedly stiffen their hand and wrist (without moving their fingers) between relaxed and completely stiff states, coordinating their change of stiffness with the cursor, relaxing when it was in the base of a trough and stiffening maximally when the cursor was at the top of the sinus peaks. We recorded the EMG from the two flexor (*E*_*f*1_, *E*_*f*2_) and two extensors (*E*_*e*1_, *E*_*e*2_) through the calibration session. According to previous studies (like Tee et al., 2010), we assumed a linear joint-muscle model where the finger movement torque (*τ*) is a consequence of the muscle tension and is correlated with the EMG, acting on the joint through a moment arm. As the participants do not move their wrist or fingers, the torque output is zero through the calibration period. Overall, these assumptions can be written as

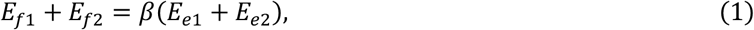

where *β* represents the linear weights that include the moment arm and the calibration constant between the EMG and muscle tension. *β* was calculated using the root mean square fit on the flexor and extensor EMGs (after a third-order Butterworth filter of cutoff frequency 8 Hz) through the calibration session.

With the *β*, one can subsequently estimate the co-activation at any time instance as

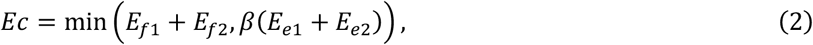

where min() represents the minimum function. The joint angle of the sixth finger (*θ*) was then decided as

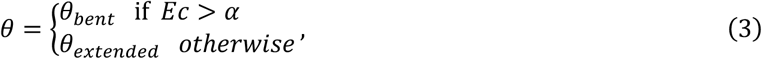

where *θ_bent_* and *θ_extended_* are the bent and extended positions of the finger. *α* was adjusted for each participant by the experimenter during the calibration session to determine the minimum *Ec* that can be recognized as the co-activation.

### Experimental procedure

#### Habituation task

After adjusting the control parameters in the above calibration, participants performed a training task to bend the fingers of the left hand (Figure 1B). The robotic finger was attached on the side of the little finger of the participant’s left hand, and the six-fingered glove was worn over it. Markers were attached to the glove to track the movement of each finger.

Participants were presented with music through a pair of headphones and were required to bend one of the six fingers in a pseudo-random order, as indicated by visual cues. The visual cues were presented aligned with the beats of the music. The participant worked in one of two habituation tasks:

##### Task 1

Ten participants performed a finger bending task synchronized with the music beats. The participants sat on a chair in front of a desk and put their elbows on the desk with their palms facing toward them. Light emitting diodes (LEDs) were fixed to the tip of the fingers of the glove worn by the participants. The LEDs lighted up to indicate which finger to be bent (see Figure 1B left).

##### Task 2

Eight participants performed a finger taping task synchronized with the music beats. Piano-like keys were displayed on a tablet PC on the desk placed in front of the participants, and one of the keys lighted up to tell which finger to be bent. The participants were asked to hold their fingers over the keys, one above each, and touch the keys on the display by bending the corresponding finger (see Figure 1B right).

Participants practiced the above tasks under two conditions: *controllable* and *random*. In the controllable condition, the robotic finger was linked to their EMG and could be manipulated voluntarily. In the random condition, the robotic finger was flexed at a predetermined random time and could not be controlled by the participant. The participants were however asked to still co-contract the wrist muscles, as with the controllable condition, when they wanted to bend the robotic finger. The order of the two conditions was randomized across the participants.

All participants were provided with haptic feedback with the stimulus pin in the controlled conditions. In the random conditions, eight participants were provided with haptic feedback using the stimulus pin, which was however synchronized with the finger movement. The remaining participants worked without the stimulus pin in the random condition. We initially planned this procedure also to examine the role of haptic feedback in the process of embodiment. However, subsequently we decided to combine the results from the two random patterns because participants reported haptic feedback due to the shifting of the weight of the finger when it moved, even when the stimulus pin was absent.

#### Pre- and post-habituation tests

We utilized two behavioral tests to examine the changes in perception caused by the training task. The tests were both before (pre) and after (post) training and were performed with the EMG sensors and the robotic finger removed. In order to avoid loss of any induced feeling of embodiment, the participants were blindfolded after the habituation task session and instructed not to move their index fingers between the habituation task session and the test sessions.

##### Reaching test

The reaching test required the participants to sit on a chair in front of the table. They placed their left hand on the table. A tablet PC (VAIO Z VJZ13AIE) placed in front of them on the table displayed a horizontal ‘target bar’ (see Figure 3A), which the participants had to reach and touch with their left index finger, extending from the left and across the center of the screen. A cardboard box was placed on the table in front of the left hand such that it blocked the direct path of the left hand to the target bar on the tablet. The box forced the participants to curve their hand trajectory to avoid colliding with the box when they touched the target bar.

A colored square was presented on the right side of the tablet (Figure 3A). The color of the square changed randomly (red, green, blue, or black). The duration of the color presentation was 500 ms for red, green, and blue and 3 s for black. The participants were instructed to move their left hand and touch the target as soon as they observed the black color of the square.

Participants performed ten reach trials each in the pre- and post-habituation test sessions. During these touch trials, we measured the reach trajectories by tracking a marker attached to their index fingernail with MAC3D System (Motion analysis Corp.) to analyze whether and how the change in the features of the reach trajectories can be attributed to changes in the perceived width the hand.

##### Finger localization test

The finger localization test required the participants to again sit on a chair in front of a table. They inserted their left hand between an open box top and the table surface while keeping the palm down and the fingers near each other so that their hand and fingers were not visible to the participants. A tablet PC was placed vertically, with the screen facing the participants, behind but adjacent to the box such that only part of the screen was above the plate was visible to the participants (Figure 3B).

The participant was presented a vertical target bar on the tablet screen, such that the part of the target above the box was visible to the participant, while the part below the box’s top was not. The participant was instructed to touch the target line below the plate (without their hand being visible) with either their index or little finger. The name of the finger to be utilized was mentioned besides the target line. The participants touched ten times with either finger, both in the pre- and post-habituation tests sessions. The index and little finger touch locations were recorded and analyzed for differences across conditions. The lateral distance between the target line and the position touched by the instructed finger defined the ‘drift’ of the finger.

#### Embodiment questionnaires

The questionnaire answers were recorded post-habituation by asking the participants to rate seven items written in Japanese on a seven-level Likert scale ranging from 0 (I did not feel that at all) to 7 (It felt extremely strong). The items of the questionnaire (translated to English) were:

Q1: During the training task, the artificial finger moved when I wanted to move it.
Q2: During the training task, the 6-fingered hand began to look normal.
Q3: During the training task, moving the artificial finger was as easy as moving the actual finger.
Q4: During the training task, I felt that the artificial finger was part of my hand.
Q5: During the training task, I felt that I was moving the artificial finger by myself.
Q6: During the training task, I felt that the artificial finger was my finger.
Q7: My hand is (narrowed/normal/widened) after removing the artificial finger.

For Q7, ‘narrowed’ is assigned to 0 and ‘widened’ is assigned to 7.

### Data analysis

#### Reaching test

The motion tracking data were smoothed with a fourth-order Butterworth filter (7 Hz) for each marker’s trajectory using Cortex (Version 7). Then the coordinates were rotated so that the X and Y axes followed the edge of the obstacle (see Figure 3A). The peak Y-coordinate recorded in the time course of the index finger in each reach was considered as the end of the reach. We hypothesized that a change in perceived hand width would lead to a change in the obstacle avoidance, and therefore analyzed three representative measures of obstacle avoidance: the total length of each reach, the X coordinate of the end point, and the difference between the end point and start point of the trajectories on each trial (see Figure 3A). The difference of the mean and standard deviation of these three measures (between the pre- and post-habituation tests sessions) gave us a measure of obstacle avoidance for each participant defined as 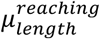 and 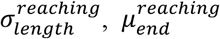 and 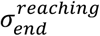, and 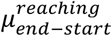 and 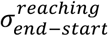, respectively.

#### Finger localization test

We calculated the finger drift by subtracting the horizontal positions touched by the instructed finger from that of the target line (see Figure 3B). The change in the mean and standard deviation of these values were analyzed between the pre- and post-habituation tests, which gave us a measure of spatial drift for each participant defined as 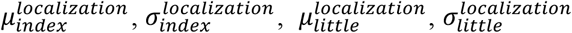. We also estimated the hand width without the thumb by 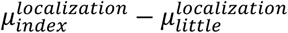.

#### Bootstrap estimation of correlation between questionnaire scores and behavior measures

To estimate correlations between questionnaire scores and behavioral measures across individual participants while suppressing influences from outliers, we applied bootstrap sampling of the questionnaire scores and behavioral measure. Bootstrap sampling was performed by the following procedures. First, a pair of questionnaire scores and behavioral measures were resampled with replacement for the same sample size (N = 18). Second, a spearman correlation value was calculated using the resampled 18-pairs of questionnaire scores and behavioral measures. Third, these steps were repeated 1000 times and the distribution of the Spearman correlation value was calculated to define a confidence interval within which 95% of the correlation values were distributed while treating both of the bottom and top 2.5% estimates as statistical outliers. We performed this procedure for every combination of seven questionnaire scores and eleven behavioral measures and evaluated 77 distributions of the correlation values between questionnaire scores and behavioral measures. Then we identified pairs of a questionnaire item and a behavior measure whose confidence interval of correlation distribution exceeded zero correlation in either of the bottom or the top edge. These pairs were considered to show significant correlation.

## Acknowledgements

The authors thank Yui Takahara for technical assistance during experiments. This research was supported by JST ERATO Grant Number JPMJER1701 (Inami JIZAI Body Project), and JSPS KAKENHI Grant Number 15K12623.

